# Programmed deep septal pacing for the diagnosis of left bundle branch capture

**DOI:** 10.1101/786665

**Authors:** Marek Jastrzębski, Paweł Moskal, Aleksander Kusiak, Agnieszka Bednarek, Tomasz Sondej, Grzegorz Kiełbasa, Adam Bednarski, Pugazhendhi Vijayaraman, Danuta Czarnecka

**Affiliations:** First Department of Cardiology, Interventional Electrocardiology and Hypertension, Jagiellonian University, Medical College, Krakow, Poland; Geisinger Heart Institute, Geisinger Commonwealth School of Medicine, Wilkes-Barre, PA, USA

**Keywords:** Left bundle branch pacing, non-selective capture, refractoriness, effective refractory period, electrocardiogram

## Abstract

**Background:** During permanent deep septal pacing, it is important to confirm left bundle branch (LBB) capture.

**Objective:** The effective refractory period (ERP) of the working myocardium is different than the ERP of the LBB; we hypothesized that it should be possible to differentiate LBB capture from septal myocardial capture using programmed extra-stimulus technique.

**Methods:** In consecutive patients undergoing pacemaker implantation who received pacing lead in a deep septal position programmed pacing was delivered from this lead. Responses to programmed pacing were categorized on the basis of QRS morphology of the extrastimuli as: myocardial (broader QRS, often slurred), selective (narrower QRS, preceded by an isoelectric interval) or non-diagnostic (unequivocal change).

**Results:** Programmed deep septal pacing was performed 269 times in 143 patients; in every patient with the use of an 8-beat basic drive train of 600 ms and when possible also during supraventricular rhythm. Responses diagnostic for LBB capture were observed in 114 (79.7%) of patients. Selective LBB paced QRS was more often seen when premature beats were introduced during the intrinsic rhythm rather than after the basic drive train. The average septal-myocardial refractory period was significantly shorter than the LBB refractory period: 263.0±34.4 ms vs. 318.0±37.4 ms.

**Conclusions:** A novel maneuver for the diagnosis of LBB capture during deep septal pacing, was formulated, assessed and found as diagnostically valuable. This method, based on the differences in refractoriness between LBB and the septal myocardium is unique in enabling the visualization of components of the usually fused, non-selective LBB paced QRS complex.

**Graphical abstract:** 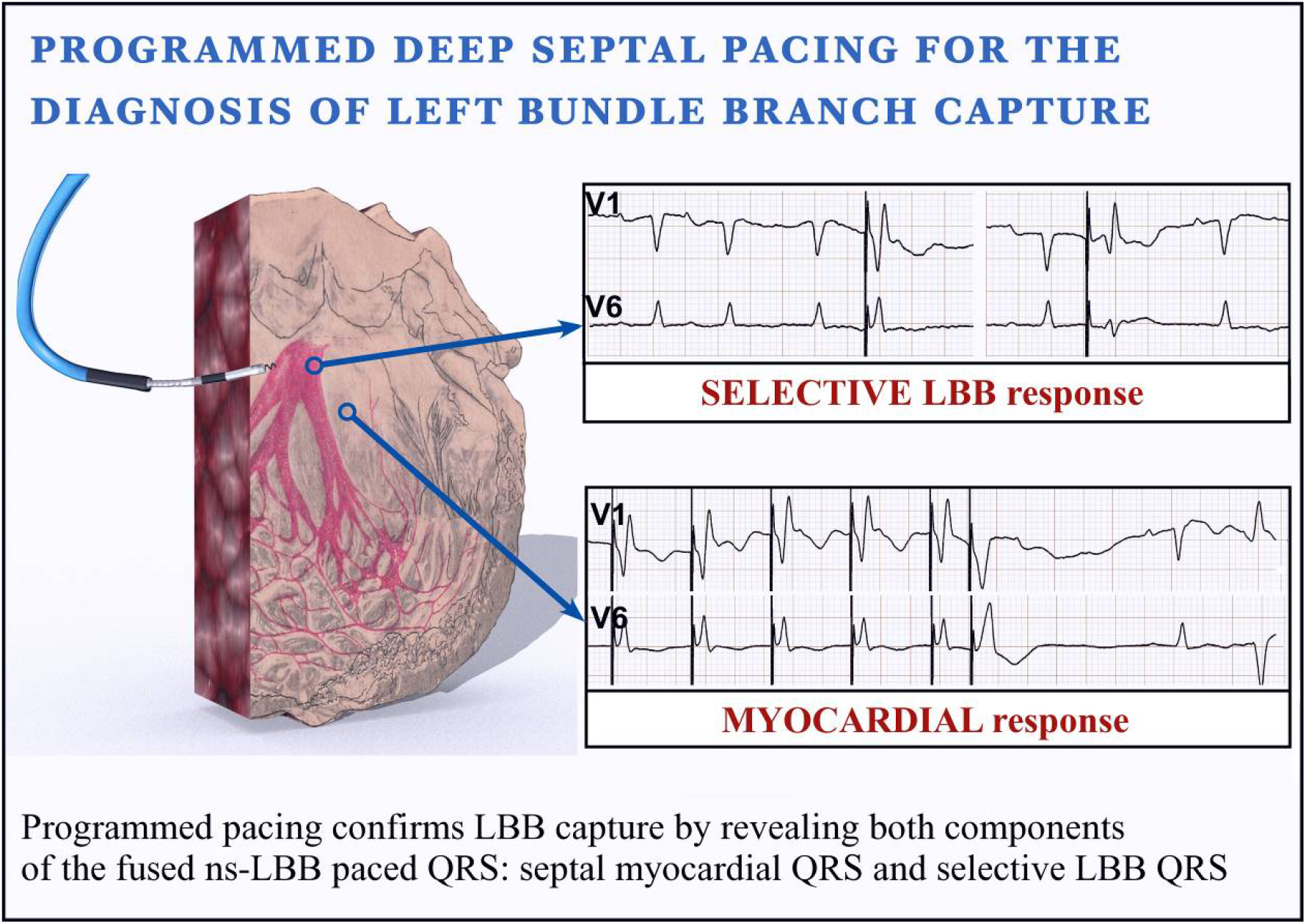

## Introduction

Permanent deep septal pacing with direct capture of the left bundle branch (LBB) is a new promising pacing option with a potential application for both bradyarrhythmias and heart failure treatment.^1–6^ During deep septal pacing it is important to ensure that the LBB or its proximal fascicles are truly captured. In the vast majority of deep septal pacing cases LBB capture is accompanied by the simultaneous engagement of the local septal myocardium, resulting in non-selective (ns) LBB capture as the predominant form of ventricular activation with this pacing modality. Both LBB-capture QRS and deep-septal-myocardial capture QRS are usually relatively narrow and of right bundle branch block morphology. Therefore, during electrocardiographic assessment of ns-LBB pacing it is not immediately apparent if direct LBB capture was achieved or just septal myocardial capture is present.

A similar issue is present during His bundle (HB) pacing, however, diagnosis of HB capture, in most cases, can be easily established using differential pacing output - a maneuver based on differences in capture thresholds between the His-Purkinje system (HPS) and the working myocardium.^7, 8^ During deep septal pacing this method often fails, as capture thresholds of LBB and of the adjacent working myocardium are usually very similar. Therefore, new methods / criteria for diagnosis of LBB capture are needed.

Programmed HB pacing, a technique that exploits differences in effective refractory period (ERP) between HB and septal working myocardium, was shown to be capable of providing definite diagnosis of HB capture during ns-HB pacing by visualizing components of the fused paced QRS complex, i.e. myocardial capture QRS and/or selective HB capture QRS.^9^ We hypothesized that programmed pacing will be able to provide definitive diagnosis of LBB capture in a similar fashion; this was never investigated before.

## Aim

To analyze responses observed during programmed deep septal pacing in regard to the diagnosis of LBB capture.

## Methods

This was a single-center prospective study: consecutive patients who received His-Purkinje system pacing device were screened and patients with pacing lead deployed deep intraseptally, in order to capture LBB, were included. To achieve deep septal pacing, a thin (4.1 F), active helix, screw-in pacing lead (model 3830, Medtronic, USA) was positioned with the help of a delivery sheath (Medtronic models C315His, C304XL and C315S10 - depending on the heart anatomy). The HB potential (if recorded) or the most superior aspect of the tricuspid annulus, appraised electrophysiologically and fluoroscopically, were used as anatomical landmarks. We aimed at any basal to mid-interventricular septal site where proper deep septal lead deployment was possible. Lead behaviors during deep septal deployment, characterized by us elsewhere,^10^ were used to guide lead positioning and fixation. The area located approximately 2 – 3 cm apically from the distal His bundle site was targeted; preferentially characterized by paced QRS morphology in V1 showing notch near the S wave nadir (“W” morphology) and/ or being slightly narrower than the paced QRS from neighboring sites and with normal axis in the frontal plane (R in lead I, Rs in lead II and rS in lead III). Lead deployment was performed under fluoroscopic and ECG guidance. Constant or interrupted pacing from the lead was delivered in order to monitor change in the paced QRS morphology during screwing-in. We aimed to obtain paced QRS with an r’ deflection in lead V1, record LBB potential and/or observe evident QRS narrowing as compared to the initial right ventricular septal paced QRS. If after 5-8 lead turns a typical progressive change of paced QRS morphology was not observed or early strong counterclockwise torque build-up in the lead was present,^10^ the lead was repositioned. This implantation technique was similar to the recently described approach to LBB pacing by Huang et al.^4^ but also to the left septal pacing method developed by Mafi-Rad at al.^5^

Programmed pacing was performed when the final deep intraseptal lead position was reached. The pacing output was set at 2 times the capture threshold and the lead was connected to an universal heart stimulator (UHS 3000, Biotronik, Germany) in a unipolar pacing configuration. Premature beats were introduced after an 8-beat basic drive train of 600 ms and also during the intrinsic rhythm (when present). The coupling interval was decreased in 10 ms steps, starting from 400 - 450 ms (or longer if the QRS of the first premature stimulus was not identical as the QRS of the drive train), until the complete loss of capture. Responses to programmed pacing were analyzed with the use of an electrophysiology system (Lab System Pro, Boston Scientific, USA) using sweep speed of 50 – 100 mm/s and categorized according to the principle delineated below. Theoretically, when the septal-myocardial ERP is shorter than the LBB ERP, the first extra-stimulus delivered at a coupling interval shorter than the LBB ERP should result in a QRS widening revealing QRS morphology of pure myocardial capture; while in cases where the septal-myocardial ERP is longer than the LBB ERP, the first extra-stimulus with a coupling interval shorter that septal-myocardial ERP should be followed by selective LBB paced QRS complex. Both responses would be diagnostic of non-selective pacing and, consequently, would confirm that LBB capture was achieved.

Changes in the paced QRS morphology from ns-LBB into selective-LBB or myocardial QRS during increase/decrease in pacing output (differential pacing output maneuver) were analyzed in every patient. The QRS complex morphologies obtained during this maneuver were compared (pairwise) with QRS complexes obtained with programmed pacing.

All patients gave written informed consent for participation in this study and the Institutional Bioethical Committee approved the study protocol.

## Results

### Population

Out of 412 consecutive patients that underwent His-Purkinje system pacing device implantation within one year, 163 patients received deep septal pacing lead, 143 of which had programmed pacing properly performed at the time of implantation and were included in the current study; clinical and pacing related characteristics of these patients are presented in Table 1.

**Table 1.** Basic clinical characteristics of the studied group (n = 143)

|  |  |
| --- | --- |
| Age [years] | 76.1±11.8 |
| Male gender | 77 (53.8%) |
| Pacing indication [n]: |  |
| 1. Sick sinus syndrome | 30 (21.0%) |
| • Atrioventricular block | 65 (45.4%) |
| • Atrial fibrillation with bradycardia | 20 (14.0%) |
| • Heart failure | 28 (19.6%) |
| Comorbidities [n]: |  |
| • Heart failure | 58 (40.8%) |
| • Coronary heart disease | 55 (38.5%) |
| • Diabetes mellitus | 61 (42.7%) |
| • Hypertension | 122 (85.3%) |
| • Severe valvular disease | 19 (13.5%) |
| LV ejection fraction [%] | 50.9±14.3 |
| LV end-diastolic dimension [mm] | 50.5 ±8.2 |
| Native QRS duration [ms] | 127.5±30.6 |
| Paced QRS duration*[ms] | 111.9 ±15.1 |
| Ventricular sensing [mV] | 9.0 ±5.1 |
| Capture threshold [V] | 0.6 ± 0.3 |

HB – His bundle, ns – non-selective, LV – left ventricular, HV – His-ventricle
\* Measured from real QRS onset rather than from pacing spike

### Programmed pacing

Programmed deep septal pacing was performed 269 times: 143 times with the use of an 8-beat basic drive train of 600 ms and when possible also during intrinsic supraventricular rhythm (126 times).

Responses observed during programmed LBB pacing were categorized as follows:

1. Diagnostic response type 1 (“myocardial”, Figure 1, left panel): change of paced QRS into a myocardial-only paced QRS, that is evidently broader, often with slur/notch/plateau instead of a pointy R wave peak and/or with obvious change in amplitude/polarity in several leads.
2. Diagnostic response type 2 (“selective LBB”, Figure 1, right panel): change of paced QRS morphology to a selective QRS of right bundle branch morphology preceded by a latency interval.
3. Non-diagnostic response: recognized when neither diagnostic response type 1 nor type 2 is present. During non-diagnostic response progressive QRS prolongation and only minor amplitude change are observed when the extrastimuli encroach on the relative refractory period of the working myocardium (final 1 - 3 coupling intervals before ERP is met).

**Figure 1.**
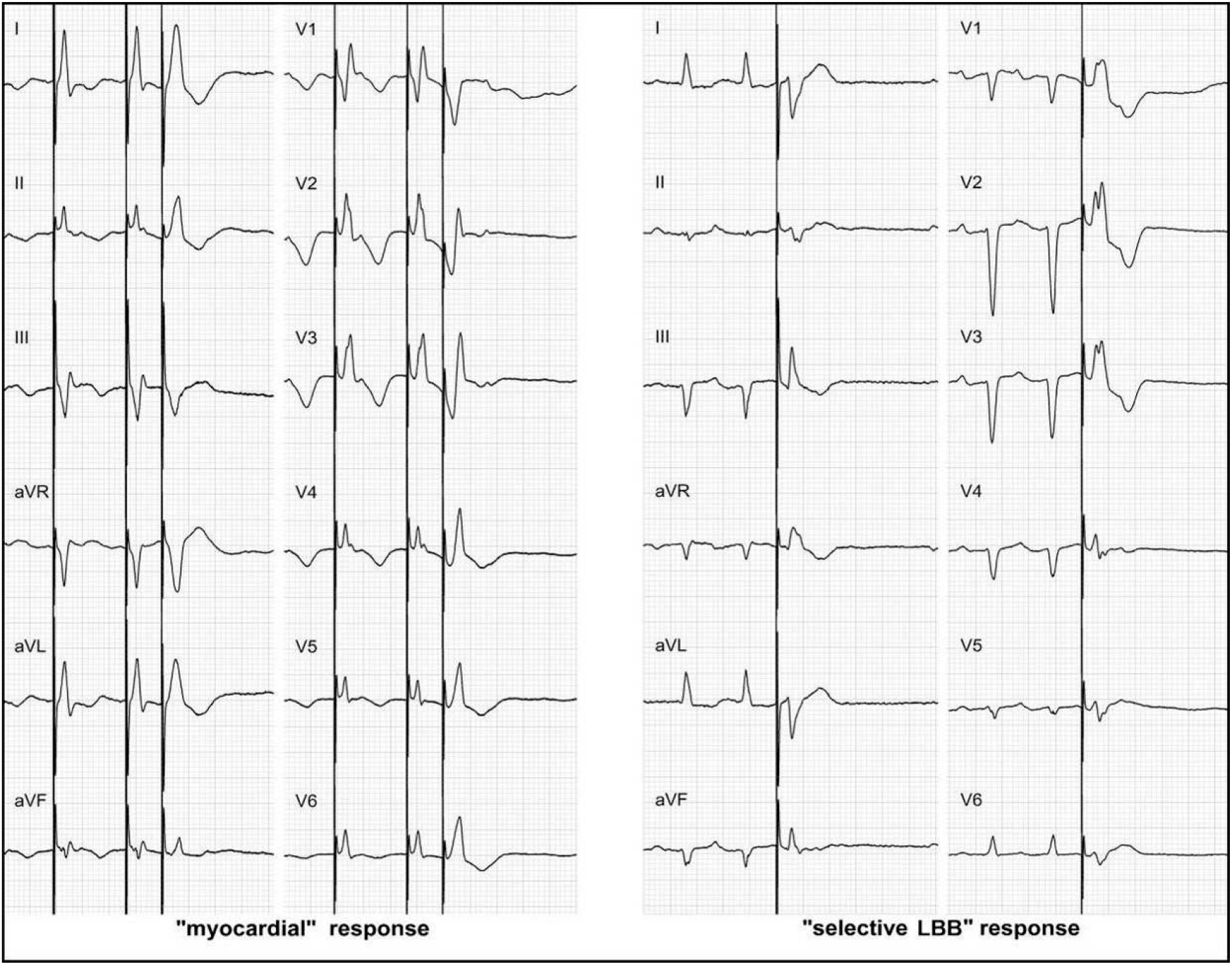
Two types of diagnostic responses to programmed deep septal pacing observed in the same patient. Left panel: extrastimulus delivered after a basic drive train of 600 ms results in a myocardial response. Note the loss of right bundle branch pattern in lead V1, QRS prolongation and loss of pointy R wave peak - both most evident in leads I, II and aVR. Right panel: extrastimulus delivered during intrinsic supraventricular rhythm results in a selective left bundle branch (LBB) capture. Note isoelectric interval before QRS, augmentation of right bundle branch morphology in lead V1 and opposite polarity of QRS in leads III and V6 as compared leads to the ‘myocardial’ response. Paper speed 25 mm/s.

A total of 114 (79.7%) patients showed diagnostic response; “selective” response was observed 44 patients while “myocardial” response in 107 patients. Both myocardial and selective responses, i.e. visualization of both components of the fused ns-LBB QRS, was possible in 37 (25.9%) of patients. In all these cases the selective LBB paced QRS and selective myocardial paced QRS were compatible with each other - with regard to the morphology of the ns-LBB paced QRS (Figure 2). Selective response was much more often seen when premature beats were introduced during the intrinsic rhythm rather than after the basic drive train, 43 (30.1%) vs. 2 (1.4%), respectively. While myocardial response was more often seen when premature beats were introduced after the basic drive train 104 (72.7%) vs 33 (23.1%), respectively.

**Figure 2.**
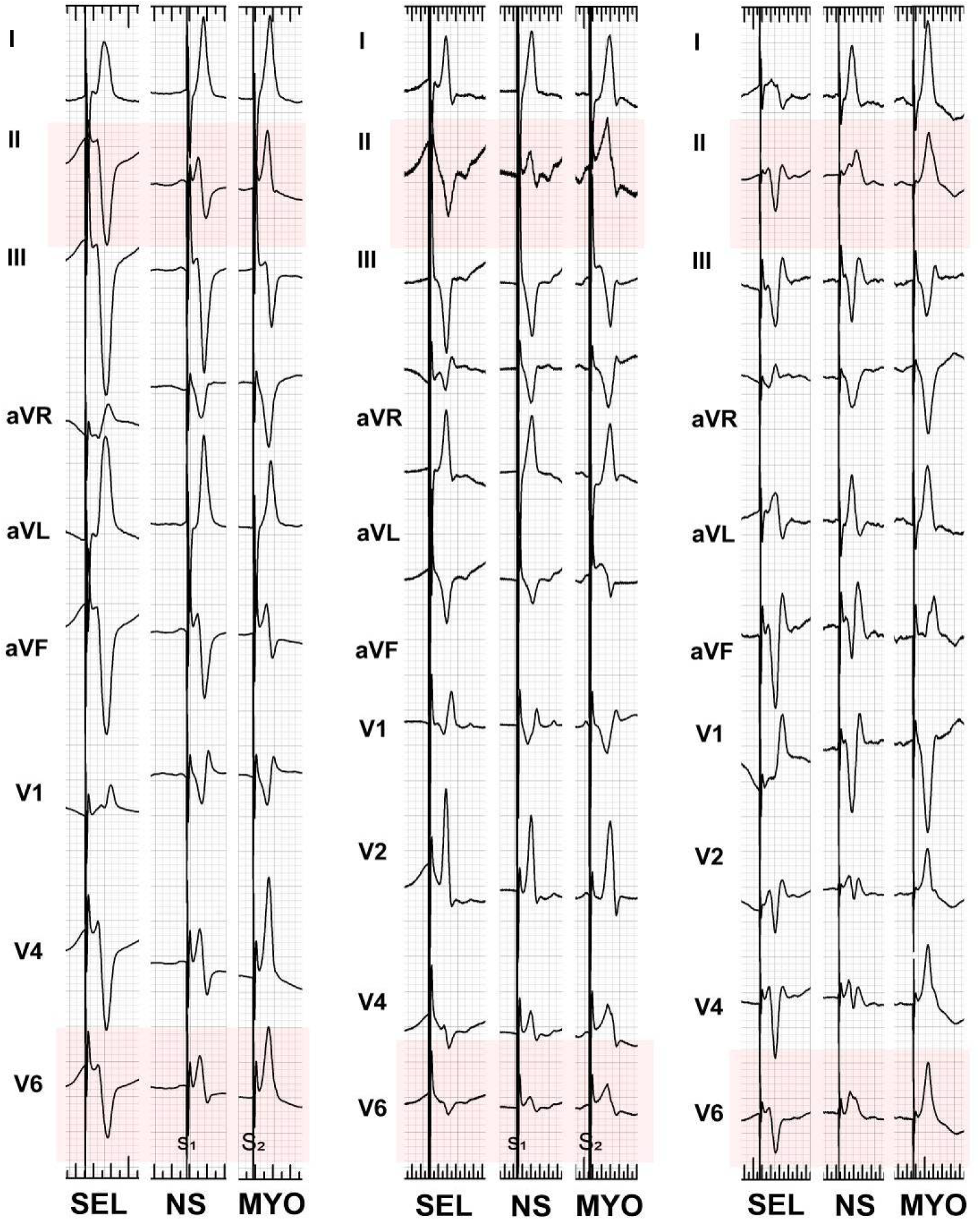
Three cases illustrating that programmed deep septal pacing can visualize both components of the non-selective (NS) left bundle branch paced QRS – that are morphologically compatible. The NS left bundle branch QRS results from fusion of the selective (SEL) left bundle branch QRS and the myocardial (MYO) only paced QRS. This can be best appreciated analyzing QRS in leads II and V6. Note QRS changes typical for loss of LBB capture: inferior axis shit and increase in QRS amplitude in precordial leads. Paper speed 25 mm/s.

The average septal-myocardial refractory period was significantly shorter than the LBB refractory period: 263.0±34.4 ms vs. 318.0±37.4 ms, p < 0.01 and 293.1±56.5 ms vs. 355.5±69.8 ms, p < 0.0001, when assessed with the 600 ms drive train or with extrastimuli delivered during supraventricular rhythm, respectively.

### Differential pacing output maneuver

QRS morphology change during decreasing/increasing pacing output was observed in 27 (18.9%) patients. In 22 (15.4%) cases, selective QRS was revealed (myocardial threshold > LBB threshold) while in 5 (3.5%) cases, myocardial QRS was revealed (myocardial threshold < LBB threshold). In all these cases, differences in capture thresholds was small, and, consequently, change of QRS morphology was observed briefly - just before complete loss of capture. In all patients in whom LBB capture was confirmed by the differential pacing output the programmed pacing also provided diagnostic response; the QRS complexes obtained by both techniques were found compatible in all cases.

## Discussion

The major finding of the current study is that programmed deep septal pacing could reveal components of the fused paced QRS complex and thus, confirm LBB capture in the majority of deep septal pacing cases. This was possible due to the differences in refractoriness and local activation times between HPS and working myocardium.

### Myocardial QRS response during programmed pacing

The current study shows that during non-selective pacing extrastimuli with short coupling interval (usually < 300 ms) results in changes of paced QRS morphology due to the selective myocardial capture i.e. loss of HPS capture. Such a QRS response was determined by ERP of the HPS being longer than ERP of the working myocardium.

Careful analysis of paced QRS morphology is crucial for the diagnosis of pure myocardial capture; sometimes changes from ns-LBB to myocardial QRS are very obvious, but at times relatively subtle or gradual. Moreover, the spectrum of myocardial QRS morphologies is wider during the loss of LBB capture than what is seen during the loss of HB capture. This is determined by a wider spectrum of pacing lead positions both in superior-inferior (QRS axis) and basal-apical (precordial transition) orientation and also with regard to the depth of penetration into the interventricular septum (left or right bundle type configuration). Our current observations suggest that change from ns-LBB to myocardial QRS usually results in a rightward (inferior) axis shift, stronger anterior forces (higher amplitude in precordial leads V2-V5), loss/decrease in amplitude of r′ in V1 (QS configuration), longer global QRS duration, delayed R wave peak in V6, notches and rounded/slurred R wave peaks (Figures 1, 2, 4 and 5). These last observations are analogous to what we have reported for the right ventricular septal myocardial QRS during loss of HB capture.^11^

**Figure 3.**
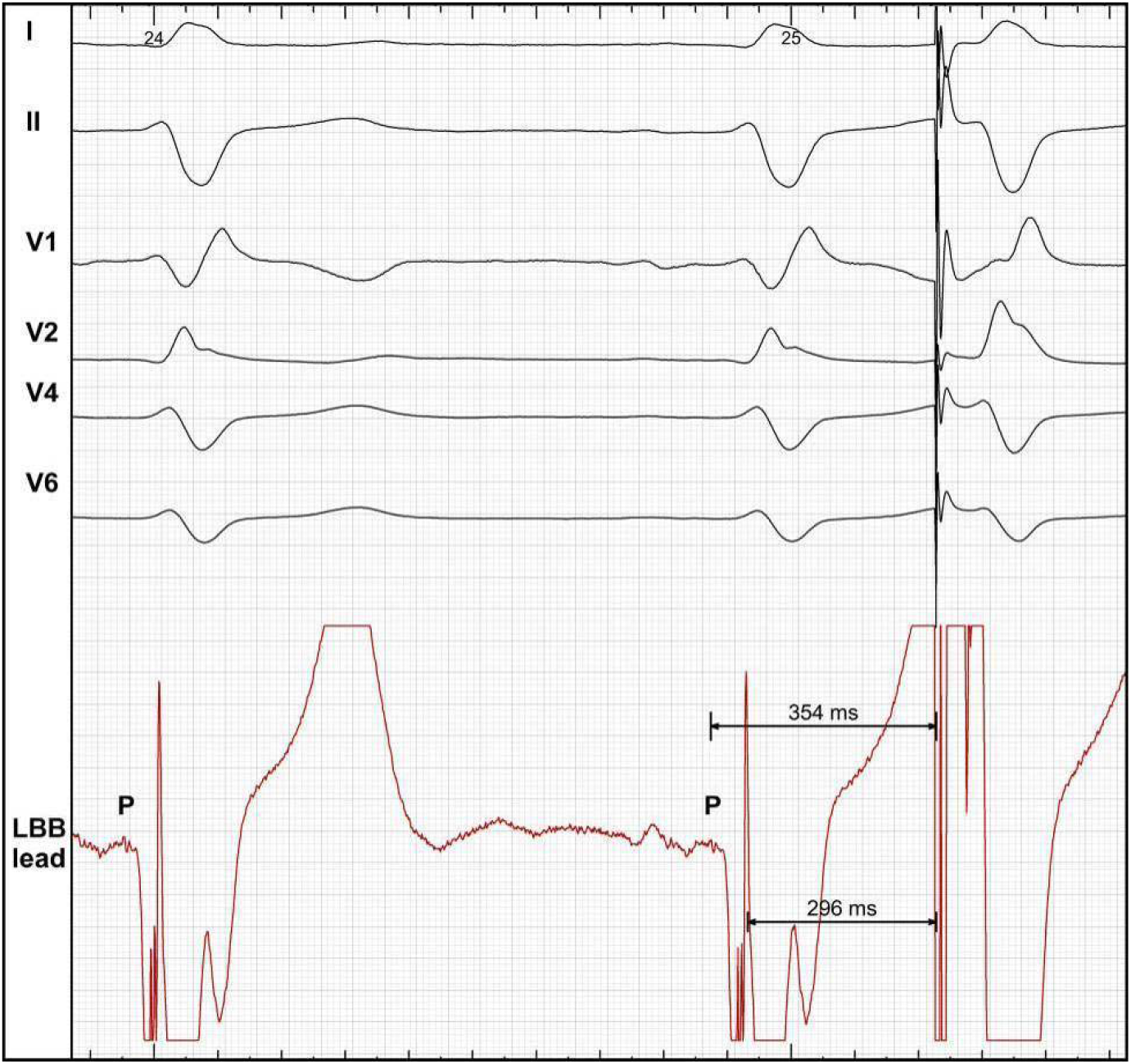
During native supraventricular rhythm with intact conduction in the left bundle branch (LBB), the LBB is always depolarized earlier than the adjacent myocardium. Consequently, when an extrastimulus is delivered during supraventricular rhythm the LBB has more time to recover (354 ms in the current example) and might be already excitable while the working myocardium has less time to recover (296 ms in the current example) and might be still refractory. This phenomenon enables to obtain selective QRS during programmed pacing even in cases when effective refractory period of LBB is longer than the myocardial effective refractory period. Note the right bundle branch block during intrinsic conduction that augments the difference in local activation times (LBB vs. adjacent myocardium). P - LBB potential.

**Figure 4.**
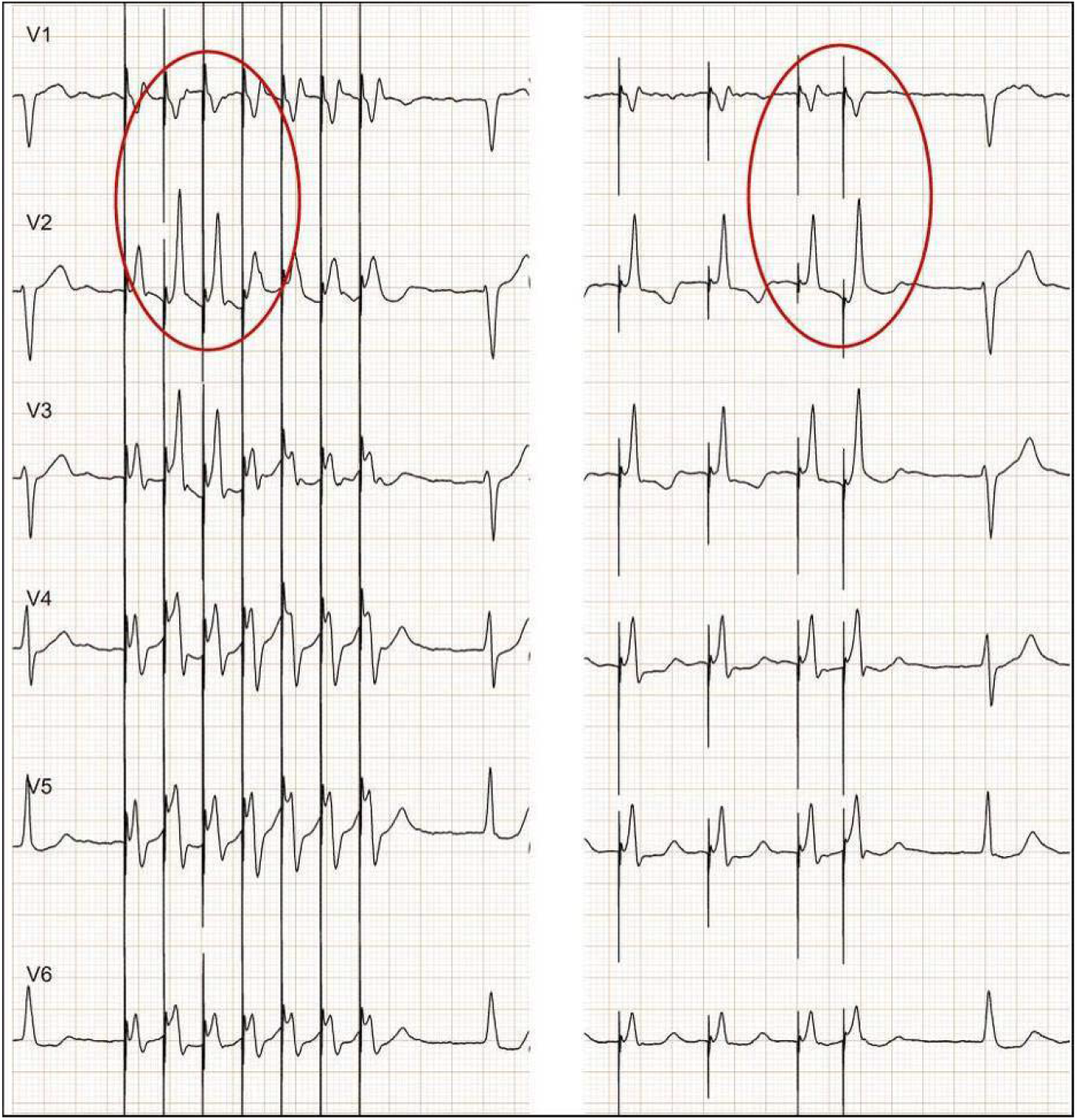
Burst pacing (left panel) with cycle length close to the effective refractory period of the left bundle branch (LBB) immediately provides the answer that LBB capture was achieved. With loss of LBB capture the r′ in V1 disappears and the prominent anterior forces emerge - this morphology was identical as myocardial QRS obtained during programmed deep septal pacing (right panel) in the same patient.

**Figure 5.**
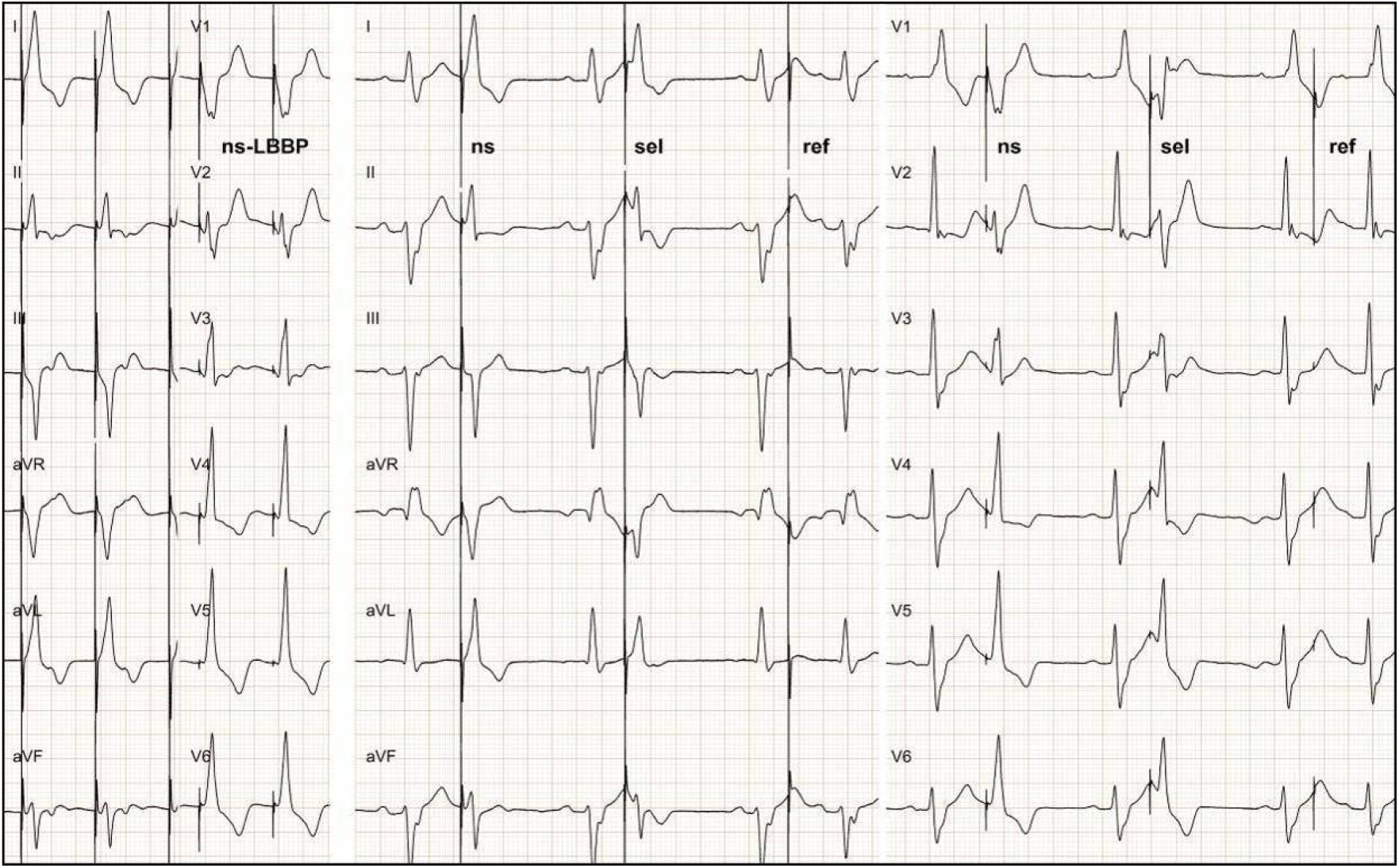
During deep septal pacing a relatively wide QRS and without r′ in V1 was obtained (left panel) in this case. Uncertainty as to the left bundle branch (LBB) capture were addressed during follow-up by temporary reprogramming to asynchronous mode (VOO 45 bpm; right panel). This promptly resulted in visualization of selective (sel) LBB paced QRS - occurring always with critical coupling of the pacing spikes. Consequently, non-selective (ns) LBB capture was diagnosed and the unusually large myocardial component in the ns-LBB QRS was attributed to the relatively shallow lead penetration into the interventricular septum. Note that the first pacing stimulus (marked “ns”) with longer coupling interval results in QRS identical as during regular pacing (marked “ns-LBBP”), while the last stimulus (marked “ref”) results in non-capture due to coupling interval shorter that effective refractory period of both LBB and working myocardium. Paper speed 25 mm/s.

When analyzing deep-septal paced QRS morphology additional data might be useful to diagnose change from non-selective LBB capture into myocardial QRS. Firstly, the QRS axis of the myocardial QRS is determined by the pacing lead position while the selective QRS axis depends solely on the HPS activation. Therefore, the more superior location of the pacing lead on the septum the more positive QRS in II, III aVF is to be expected with loss of HPS capture. Secondly, the myocardial QRS morphology should be compatible with the non-selective and selective QRS morphologies in a particular case - according to the rules of QRS fusion. For example, when the ns-QRS in lead II has RS configuration and selective QRS has rS/QS configuration (due to the often seen preferential activation of the posterior fascicle), then the myocardial QRS must have R/Rs configuration in this lead (Figures 1 and 2).

Interestingly, the myocardial response during programmed LBB pacing was observed less frequently than during programmed HB pacing (74.8% vs. 100%).^9^ This might be related to the lower percentage of cases with HPS capture during deep septal pacing as compared to HB pacing. However, a non-diagnostic response was also observed in cases with a *bona fide* LBB capture (non-diagnostic response with extrastimuli delivered during intrinsic rhythm yet diagnostic response with extrastimuli delivered after a drive train - and *vice versa*). Perhaps the occasional lack of myocardial response during ns-LBB capture can be also explained by a smaller difference between ERPs of HPS and adjacent myocardium during LBB pacing as compared to HB pacing, 55 ms vs. 81.9 ms, respectively.^9^ This gives a smaller ‘opportunity’ for the extrastimuli to capture only the adjacent myocardium. It is also possible that in some cases the myocardial QRS and ns-LBB QRS are too similar and the loss of HPS capture is difficult/impossible to establish by analysis of changes of QRS morphology. Perhaps in some cases even myocardial capture might result in a relatively fast engagement of the septal HPS limiting the difference between myocardial paced QRS and ns-LBB paced QRS.

### Selective LBB QRS response during programmed pacing

A paced QRS with an isoelectric interval preceding it is considered an unmistakable diagnostic hallmark of selective HPS capture. Selective LBB QRS during programmed pacing was almost exclusively seen when premature stimuli were delivered during intrinsic rhythm. This is because during native supraventricular rhythm HPS is always depolarized earlier than the adjacent myocardium. Consequently when a HPS extrastimulus is delivered during supraventricular rhythm the HPS might be already excitable while the working myocardium is still refractory. This phenomenon enables to obtain selective QRS during programmed pacing even in cases when ERP of HPS is longer than myocardial ERP.^9^ In contrast to HB pacing a wide spectrum of selective LBB QRS morphologies is observed. It seems that with proximal capture of the LBB there is a RS configuration in lead II and rS/QS in lead III; while with inferior lead positions selective QRS imitates a left anterior fascicular block configuration (rS/QS in II, III, aVF) and with superior lead position positive QRS in II, II, aVF is observed (suggestive of preferential activation of the left anterior fascicle).

During HB pacing a selective response was observed by us in 58.8% of cases while in the current study, during LBB pacing, in only 30.8 % of cases. This is may be explained by the fact that the HB is activated 20 ms earlier than LBB and the basal septum on the right side is depolarized later than the mid-interventricular septum on the left side. Consequently, the difference in local activation times (HPS vs. adjacent myocardium) at the pacing lead site is less during LBB pacing than during HB pacing (Figure 3).

Importantly, myocardial and non-selective QRS from the deep septal (left ventricular subendocardial) site is often characterized by a considerable latency/initial isoelectric component, prominent especially in the precordial leads V2-V6. Care should be taken not to mistaken this with a latency interval of selective capture; observation of a small pseudodelta in other leads (V1, I, II, III, aVF) are helpful in the differentiation.

### Clinical translation

It is our current device implantation policy to attempt His-Purkinje system pacing in all patients with an indication for permanent cardiac pacing. Regardless of the final implemented pacing mode (HB or LBB), confirmation of HPS capture is always attempted with the same methods: programmed pacing and differential pacing output maneuvers. In our experience these maneuvers can be easily incorporated into a standard implantation protocol. However, this requires the pacing lead to be connected to an electrophysiological system with constant endocardial and surface 12-lead ECG signal acquisition and recording. Importantly, during LBB pacing quite different responses with these maneuvers are observed than during HB pacing. During LBB pacing equal capture thresholds are common (75% vs. 8-10% when compared to HB pacing) resulting in a much lower diagnostic yield of the differential pacing output maneuver. Consequently, when LBB pacing is implemented, we rely more on programmed pacing technique to confirm HPS capture. To further exploit differences in refractoriness between the HPS and the working myocardium, we occasionally use burst pacing (Figure 4) and double or triple extrastimuli with a long-short sequence, which facilitates pure myocardial capture. During the follow-up, a slow asynchronous pacing - resulting in variable coupling intervals with intrinsic QRS complexes - usually promptly provides evidence of LBB capture (Figure 5).

Programmed deep septal pacing shows that LBB capture can be present despite a relatively wide paced QRS complex, even without r’ in lead V1 (Figure 5). This suggests that LBB capture is achievable already at a substantial distance from the LBB and that from practical point of view, LBB capture is not an ‘all or nothing’ phenomenon. On the contrary, the contribution of the LBB capture and myocardial capture to the ns-LBB QRS can vary depending on the depth of penetration of the pacing lead into the interventricular septum - the more shallow the lead position the bigger contribution of direct myocardial capture. This observation corroborates our earlier report that with constant pacing from the pacing lead, during deployment into the interventricular septum (“pacemapping while screwing-in technique”), the QRS transition from a right ventricular paced QRS to a ns-LBB QRS is undeniably continuous, rather than sudden (Figure 6).

**Figure 6.**
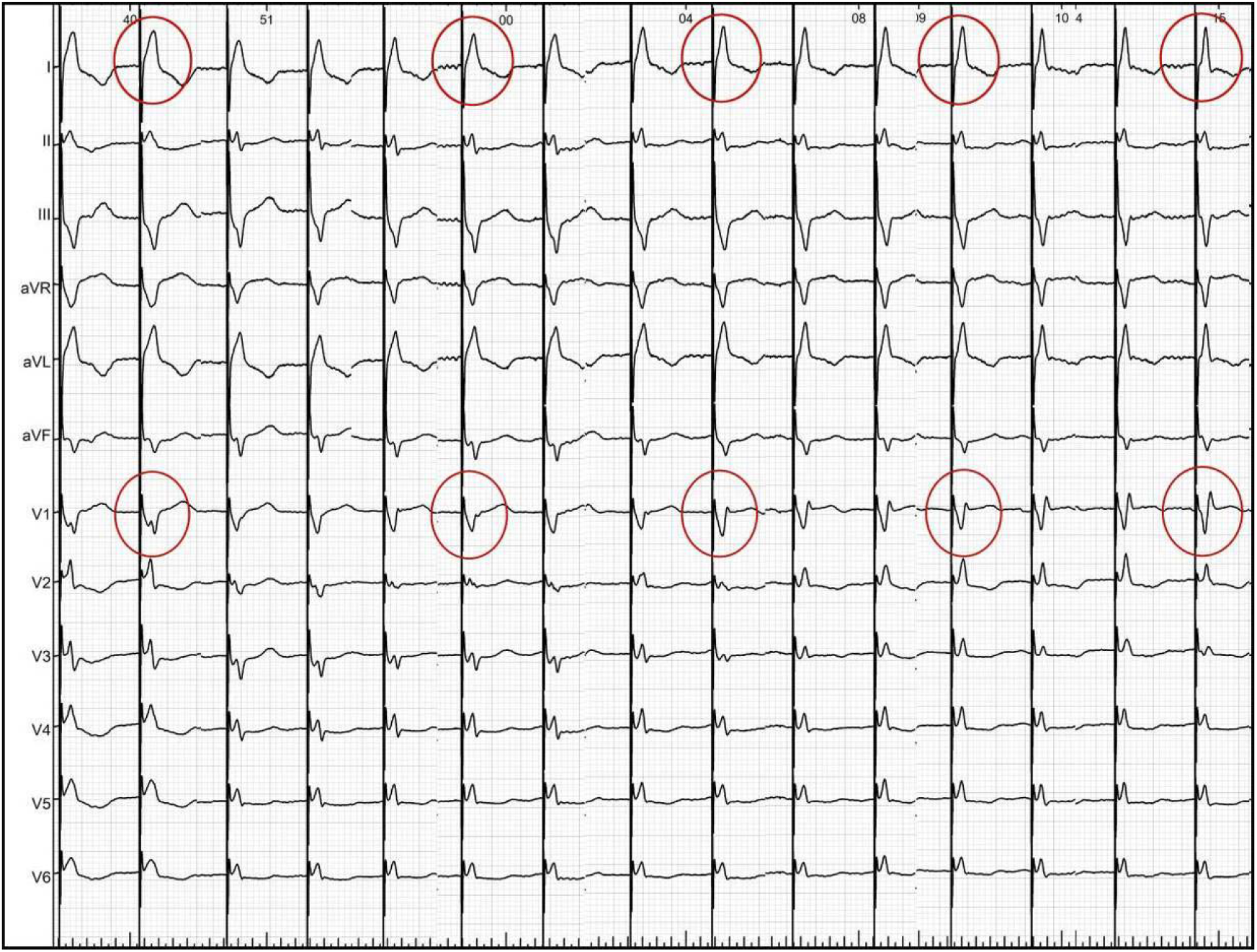
This is a collage of QRS complexes taken from a continuous ECG recorded as the lead was progressively deeper deployed into the interventricular septum without interruption of pacing (“pacemapping during screwing-in”). During this 25-seconds-long ECG there was a continuous, smooth QRS transition (circles) from the wide, right ventricular paced QRS (on the left) to the narrow, sharp QRS with r′ in V1 of the left bundle branch capture morphology (on the right). Note the transition of the V1 notch to the end of QRS and then transformation of it into r’ of growing amplitude. Despite careful analysis of all transitional QRS complexes there was not a single moment when occurrence of LBB capture could be pinpointed.

Diagnostic responses observed during either programmed pacing or differential pacing output reassures the implanting electrophysiologist that the acute goal of the procedure was achieved. However, it is more difficult to recommend how to interpret and proceed when only a non-diagnostic response is observed. Perhaps when this is accompanied by an undesirable paced QRS characteristics (duration > 140 ms, V6 RWPT > 100 ms) and lack of LBB potential, then myocardial-only capture should be strongly suspected and the pacing lead should be repositioned or deeper fixation should be attempted.

Uncertainty as to the presence/absence of LBB capture during deep septal pacing resulted in several indirect diagnostic criteria proposed recently. These include: 1.) LBB potential recorded by the pacing lead,^2, 4^ 2.) short paced QRS duration (<130 ms),^3^ 3.) short R-wave peak time (RWPT < 75 ms) in lead V6,^12^ 4.) lack of RWPT shortening with high output pacing.^4^ However, these criteria were never validated and all have potential limitations – as delineated below. It is known that the recording of HB potential by the pacing lead does not ensure HB capture and vice-versa - lack of HB potential does not excludes HB capture. During HB pacing both QRS duration and V6 RWPT values show typical Gaussian distribution with a significant overlap between myocardial capture and ns-HB capture.^11^ It is very likely that similar situations are present during ns-LBB pacing. Shortening of the V6 RWPT with high output pacing might reflect not only HPS capture but also more rapid/earlier/wider depolarization of the working myocardium, resulting in overcoming of latency, and, consequently, better fusion of the two depolarization wavefronts. Similarly, the lack of change in RWPT duration in response to high output pacing does not ensures HPS capture as such a reaction is also seen during myocardial pacing. Therefore, until these potential criteria for LBB capture are validated and their diagnostic properties firmly established it seems prudent to rely on programmed pacing and differential pacing output - techniques with firm physiological rationale behind them.

### Limitations

Single center design could result in some bias both at the level of patient inclusion and interpretation of ECG results.

This study was performed by the operators with substantial experience in both HPS pacing and general electrophysiology; programmed pacing might be more difficult to implement as a routine diagnostic tool in the setting of an implantation laboratory without the same electrophysiological background.

Encroachment on the relative refractory period of the working myocardium makes QRS morphology change difficult to interpret (is QRS prolongation/amplitude change due to the loss of LBB capture or due to the relative refractoriness of the myocardium?). To overcome this limitation, we disregarded changes in QRS morphology that occurred only with final coupling intervals (longer by less than 30 ms than the myocardial ERP).

## Conclusions

Programmed deep septal pacing adds to the armamentarium of techniques available to the diagnosis of LBB/HPS capture both during device implantation and follow-up. Visualization of both components of the fused ns-LBB paced QRS is not only reassuring with regard to the achievement of LBB capture but also gives insights with respect to the HPS activation and myocardial activation patterns in a particular case. Familiarity with myocardial capture QRS morphology might be helpful to diagnose the loss of LBB capture during electrocardiographic follow-up. Perhaps programmed deep septal pacing technique could serve as one of the ‘gold standard’ methods to validate other criteria for LBB capture; this deserves further investigations.

